# Trapped Topoisomerase II initiates formation of *de novo* duplications *via* the nonhomologous end-joining pathway in yeast

**DOI:** 10.1101/2020.05.03.075358

**Authors:** Nicole Stantial, Anna Rogojina, Matthew Gilbertson, Yilun Sun, Hannah Miles, Samantha Shaltz, James Berger, Karin C. Nitiss, Sue Jinks-Robertson, John L. Nitiss

**Author notes:** These authors contributed equally to this work (Nicole Stantial, Anna Rogojina). Greehey Children’s Cancer Research Institute, University of Texas Health Science Center at San Antonio, San Antonio, TX 78229. PPD, Inc., Middleton, WI 53562. Laboratory of Molecular Pharmacology and Developmental Therapeutics Branch, Center for Cancer Research, National Cancer Institute, National Institutes of Health, Bethesda, MD 20892. Department of Pharmaceutical Sciences, University of Wisconsin, Madison, WI 53705.

## Abstract

Topoisomerase II (Top2) is an essential enzyme that resolves catenanes between sister chromatids as well as supercoils associated with the over- or under-winding of duplex DNA. Top2 alters DNA topology by making a double-strand break (DSB) in DNA and passing an intact duplex through the break. Each component monomer of the Top2 homodimer nicks one of the DNA strands and forms a covalent phosphotyrosyl bond with the 5’ end. Stabilization of this intermediate by chemotherapeutic drugs such as etoposide leads to persistent and potentially toxic DSBs. We describe the isolation of a yeast *top2* mutant (*top2- F1025Y,R1128G*) whose product generates a stabilized cleavage intermediate *in vitro*. In yeast cells, overexpression of the *top2- F1025Y,R1128G* allele is associated with a novel mutation signature that is characterized by *de novo* duplications of DNA sequence that depend on the nonhomologous end-joining pathway of DSB repair. Top2-associated duplications are promoted by the clean removal of the enzyme from DNA ends and are suppressed when the protein is removed as part of an oligonucleotide. *TOP2* cells treated with etoposide exhibit the same mutation signature, as do cells that over-express the wild-type protein. These results have implications for genome evolution and are relevant to the clinical use of chemotherapeutic drugs that target Top2.

**SIGNIFICANCE STATEMENT:** DNA-strand separation during transcription and replication creates topological problems that are resolved by topoisomerases. These enzymes nick DNA strands to allow strand passage and then reseal the broken DNA to restore its integrity. Topoisomerase II (Top2) nicks complementary DNA strands to create double-strand break (DSBs) intermediates that can be stabilized by chemotherapeutic drugs and are toxic if not repaired. We identified a mutant form of yeast Top2 that forms stabilized cleavage intermediates in the absence of drugs. Over- expression of the mutant Top2 was associated with a unique mutation signature in which small (1-4 bp), unique segments of DNA were duplicated. These *de novo* duplications required the nonhomologous end-joining pathway of DSB repair, and their Top2-dependence has clinical and evolutionary implications.

## INTRODUCTION

Topoisomerases are enzymes that transiently cut DNA in a highly regulated fashion to carry out topological alterations. Type I topoisomerases are typically monomers that make single-strand breaks in DNA while type II enzymes are homodimers that cleave both DNA strands to create double-strand breaks (DSBs) (1-3). The ability to cut DNA is absolutely required to change DNA topology and cleavage occurs through formation of a transient, covalent phosphotyrosyl linkage with DNA. Type IA and IB enzymes create 5’- and 3’- phosphotyrosyl links, respectively, while type II enzymes form a 5’-phosphotyrosyl link. The resulting single- or double-strand break alters DNA winding using a swiveling (Type IB) or strand passage mechanism (Type IA and Type II), after which the phosphotyrosyl bond is reversed to restore the phosphodiester backbone of the DNA. While the use of a covalent enzyme-DNA intermediate makes cleavage and rejoining a relatively error-free process, any DNA breakage is potentially dangerous (4). If DNA is cut and the break persists, a DNA damage response is activated that leads to cell cycle arrest, senescence or apoptosis (5-7). Stabilization of covalent DNA-topoisomerase intermediates has been exploited to identify and develop therapeutic small molecules such as fluoroquinolone antibiotics and a variety of anti-cancer drugs (8, 9). These inhibitory molecules have also been very useful as probes for enzyme mechanism (10-12).

In addition to small molecules that trap topoisomerases on DNA, a variety of DNA structural alterations can affect cleavage and re-ligation. For type I enzymes DNA lesions that include abasic sites, nicks, base-base mismatches and base alterations can lead to enhanced DNA cleavage *in vitro* and *in vivo* (13-15). Notably, single ribonucleotides embedded in duplex DNA lead to elevated cleavage by type IB topoisomerases *in vitro* (16) and are associated with a distinctive mutation signature *in vivo* (17). In the yeast *Saccharomyces cerevisiae*, this signature is composed of deletions that remove a single unit from a tandem repeat and reflects sequential cleavage of the same DNA strand by Top1 (18-20). Type II topoisomerases can also be trapped on DNA by structural alterations, although the range of lesions appears more limited than for type I enzymes (21). Lesions that can trap eukaryotic type II topoisomerases include abasic sites (22) and mis-incorporated ribonucleotides (23). The trapping of both subunits of Top2 results in a DSB, while the trapping of only a single subunit leads to a persistent single-strand nick.

The study of cytotoxic and mutagenic mechanisms of elevated topoisomerase cleavage has been facilitated by the identification of mutants that are proficient for DNA cleavage but defective in re-ligation (24-26). While mutations that result in stabilized cleavage intermediates have been readily obtained in yeast *TOP1*, few examples have been identified for type II topoisomerases. The sole exception is a mutant form of human TOP2α (Asp48 changed to Asn) identified in a screen for mutations affecting the action of bisdioxopiperazines (27). Similar to yeast *top1* mutants, the human TOP2α^D48N^ mutant exhibits elevated DNA cleavage *in vitro* and cannot be expressed in recombination-defective (*rad52*Δ) yeast cells. We describe here a mutant yeast Top2 protein (Top2-F1025Y,R1128G; abbreviated here as Top2-FY,RG) that similarly is lethal when overexpressed in a *TOP2 rad52*Δ background and leads to elevated DNA cleavage *in vitro*. The corresponding amino acid changes, however, are in the C-terminal dimerization domain of Top2, thereby implicating this domain in the regulation of DNA cleavage and/or re-ligation by the enzyme. Top2-FY,RG overexpression elevates homologous recombination and spontaneous mutagenesis, and is associated with a distinctive mutation signature (*de novo* duplications) that is dependent on the nonhomologous end-joining (NHEJ) pathway of DSB repair. Similar *de novo* duplications were observed following treatment of WT cells with etoposide, a chemotherapeutic drug that stabilizes the covalent Top2 cleavage intermediate. Finally, we implicate WT Top2 as the source of rare *de novo* duplications observed previously in frameshift reversion assays (28). These data suggest important roles for Top2-dependent mutagenesis in genome evolution as well as in genetic stability following chemotherapy.

## RESULTS

### C-terminal *TOP2* mutations confer hypersensitivity to etoposide

We previously described mutations in yeast *TOP2* that result in hypersensitivity to different classes of Top2-targeting agents such as etoposide and mAMSA (29). Some of the drug-hypersensitive alleles were identified fortuitously using mutants constructed for other purposes while others were identified in genetic screens. In some cases, the corresponding changes were in regions of Top2 not obviously involved in enzyme-mediated cleavage or religation of DNA. One such mutant, *top2-L1052I*, was hypersensitive to mAMSA, etoposide and the fluoroquinolone CP115,953, but was only 2-3 fold more sensitive to each of these agents than a WT strain. We, therefore, carried out an additional screen following the introduction of random mutations by error-prone PCR into a DNA segment specifying amino acids 900 to 1250. The mutagenized fragment was recombined into gapped plasmid pDED1Top2 (29, 30) by cotransformation of the gapped plasmid and the mutagenized fragment into a temperature-sensitive *top2-4* strain (31). Expression of *TOP2* in strains carrying pDED1Top2 results in ∼10- fold more protein than does expression from the native *TOP2* promoter (30). Transformants were selected at 34°, which is non-permissive for the *top2-4* allele, to limit the isolation of plasmids carrying *top2* null alleles. Individual colonies were screened for sensitivity to mAMSA on solid medium and two mutants with alterations in the C-terminal part of cleavage/ligation domain of Top2 were identified. The first mutant had Arg1128 changed to Gly (R1128G) while the second (independent) mutant had the R1128G change plus an additional mutation that converted Phe1025 to Tyr (F1025Y).

The F1025Y and R1128G changes (FY and RG, respectively) were re-introduced into the pDED1Top2 plasmid individually or together by site-directed mutagenesis. Following introduction into the *top2-4* background, transformants were assessed for survival following exposure to varying concentrations of mAMSA or etoposide at 34°. *top2-4* cells carrying the pDED1Top2 or pDED1Top2-FY plasmid were insensitive to mAMSA even at a concentration of 50 µg/ml (Supplemental Figure S1A). By contrast, cells containing the *top2-RG* or *top2-FY,RG* allele were sensitive to 5 µg/ml mAMSA. *top2-4* cells carrying the pDED1Top2 plasmid were modestly sensitive to etoposide (Figure 1A) and those with the pDED1Top2-RG plasmid were more sensitive to the drug. Although over-expressing the *top2-FY* allele alone did not enhance etoposide sensitivity relative to the *TOP2* control, the double-mutant *top2-FY,RG* allele was associated with a high level of drug sensitivity. Based on drug-sensitivity profiles, only strains containing the plasmid-encoded *top2-RG* or *top2-FY,RG* allele were further analyzed.

**Figure 1.**
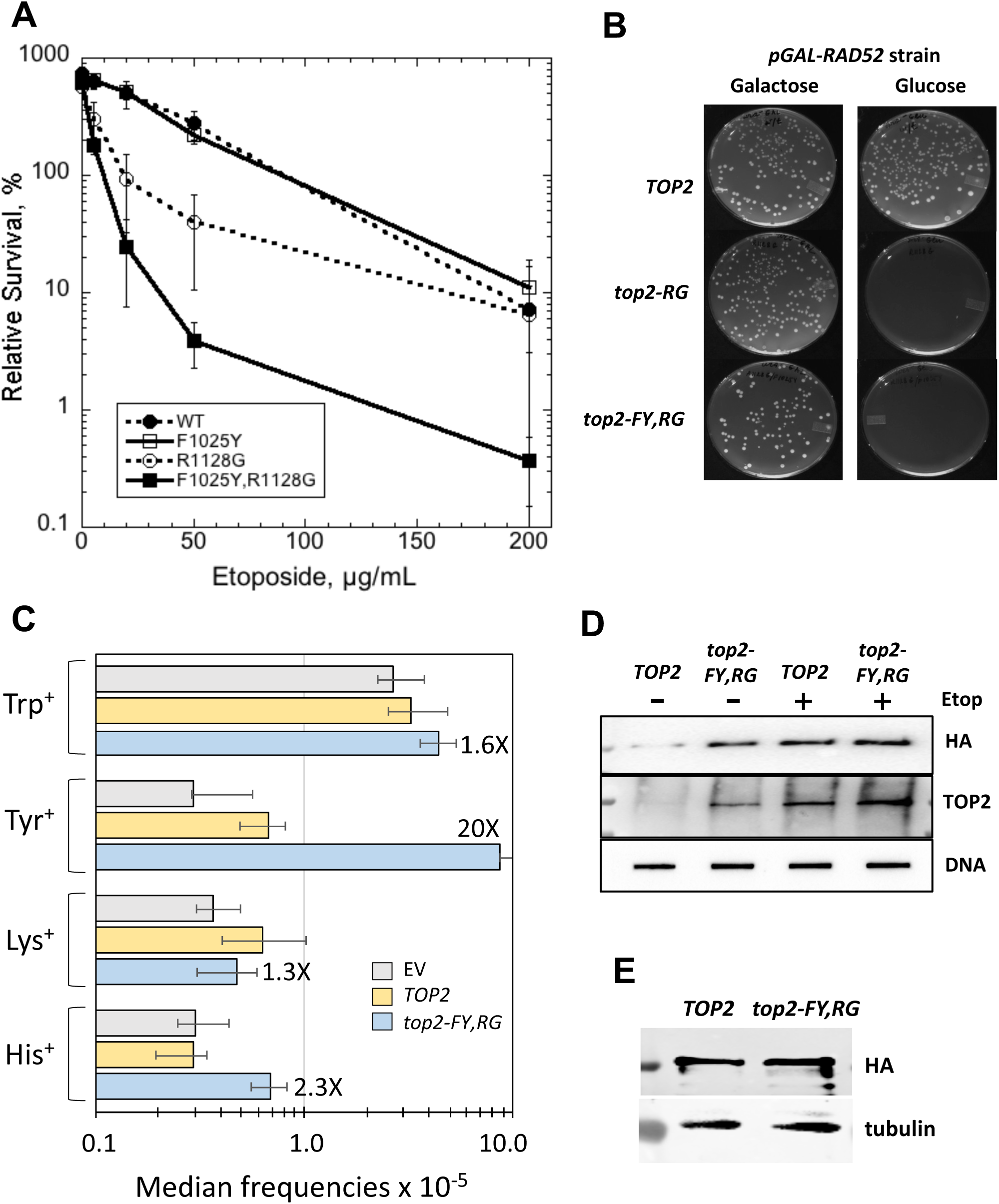
Genetic characterization of *top2* mutants. (A) The *top2-4* host strain JN362at2-4 (30) was transformed with pDED1Top2 plasmids containing the indicated alleles. Transformants were grown to mid-log phase in SC-Ura medium at 34° and the indicated concentration of etoposide was added. Incubation was continued for an additional 24 h before plating cells for survival. The survival plotted is relative to that at the time of etoposide addition. Error bars are ±SEM. (B) The pGAL-Rad52 plasmid (75) was transformed into a *top2-4 rad52*Δ background (JN332at2-4) and then cells grown in galactose medium were transformed with pDED1Top1 containing the indicated allele. Transformants were selected on uracil-deficient medium containing either glucose or galactose as a carbon source. (C) Diploid strain CG2009 was transformed with the relevant plasmid and grown selectively in SC/peptone-Ura medium prior to plating on drop-out media (SC/peptone) to select recombinants. Gray bars, EV; yellow bars, pDED1Top2; blue bars, pDED1Top2-FY,RG. Error bars are 95% confidence intervals. (D) YMM10t2-4, a *top2-4* derivative of YMM10 (35), and was transformed with the pDED1Top2 or pDED1top2-FY,RG plasmid containing an HA-tagged allele. Tranformants were grown in the presence or absence of 200 μg/ml etoposide prior to genomic DNA isolation using the yeast ICE protocol. After DNA recovery, quantitation, and digestion with miccrococcal nuclease, DNA- associated proteins were separated by SDS-PAGE and Top2 was detected using an anti-HA or anti-TOP2 antibody. Survival data following 24 h growth in the presence of mAMSA are shown in Figure S1. (E) Top2 and Top2-FY,RG levels relative to tubulin are shown.

### Expression of *top2-RG* or *top2-FY,RG* is lethal in the absence of *RAD52*

Previous drug-hypersensitive *top2* mutants were viable in a *rad52*Δ background that is unable to repair DSBs (32, 33). We were unable, however, to obtain colonies at 34° following the transformation of either the pDED1Top2-RG or pDED1Top2- FY,RG plasmid into a *top2-4 rad52*Δ strain. To more rigorously demonstrate lethality of the *top2* alleles in the absence of recombination, we introduced a plasmid containing a *pGAL-RAD52* fusion into a *top2-4 rad52*Δ background. Cells were thus phenotypically Rad52^+^ or Rad52^-^ on galactose- or glucose-containing medium, respectively. Cells transformed with pDED1Top2 grew well on both types of plates, while those carrying the mutant pDED1Top2-RG or pDED1Top2-FY,RG plasmid formed colonies only on galactose-containing medium. (Figure 1B). This effect was verified by streaking colonies obtained on galactose medium in parallel onto medium containing either glucose or galactose. Introduction of pDED1Top2-RG or pDED1Top2-FY,RG also failed to yield colonies following transformation of a *TOP2 rad54*Δ or *TOP2 rad50*Δ mutant from the yeast deletion collection. Thus, even in the presence of a WT chromosomal *TOP2* allele, over-expression of either mutant protein was lethal in the absence of recombination.

Given the requirement for homologous recombination functions for survival, we hypothesized that expression of the mutant Top2 proteins would confer a hyper-recombination phenotype. This, as well as other genetic analyses, was performed using only the *top2-FY,RG* allele since it conferred a stronger drug-sensitive phenotype. The corresponding mutant plasmid, WT pDED1Top2 or the empty vector (EV) YCp50 (34) was introduced into a diploid yeast strain carrying multiple heteroallelic markers. Recombination frequencies were measured by selecting for tryptophan, lysine, histidine or tyrosine prototrophs. Relative to the EV control, over-expression of *TOP2* from *pDED1* did not enhance recombination frequencies between any of the heteroallelic pairs examined (Figure 1C). Over-expression of the *top2-FY,RG* allele, however, resulted in 2.3-fold increase in His^+^ recombinants and a 20-fold increase in Tyr^+^ recombinants; there was no significant increase in Lys^+^ or Trp^+^ recombinants. The reason for the variable recombination effects of *top2-FY,RG* allele is unclear, but could reflect the relative distances between heteroalleles at the loci monitored, differing levels of gene expression, preferred sites of Top2 cleavage, or other features related to local chromosome structure.

### Top2-FY,RG is trapped on yeast DNA in the absence of etoposide

Top2 poisons such as etoposide interfere with DNA re-ligation catalyzed by the enzyme and create DNA damage in the form of strand breaks with covalently attached protein at the ends. Adducts formed *in vivo* can be detected using an ICE (*in vivo* complex of enzyme) assay that immunologically detects protein covalently associated with DNA isolated in a CsCl gradient (35, 36). For the ICE assay in yeast, we constructed variants of WT and double-mutant pDED1 plasmids in which the C-terminus of the Top2 protein was tagged with HA. These variants were then expressed in a *top2-4* strain with increased etoposide sensitivity due to the absence of multiple drug-efflux pumps (35). Cells expressing WT *TOP2* showed a faint Top2 signal that was greatly enhanced when cells were treated with etoposide (Figure 1D). By contrast, cells expressing the double-mutant allele showed a robust signal in the absence of etoposide that was further enhanced by drug treatment. Given the similar levels of the Top2 and Top2-FY,RG proteins (Figure 1E), we conclude that Top2-FY,RG is a self-poisoning enzyme that frequently becomes trapped on DNA in the absence of etoposide.

### Relaxation and cleavage activities of mutant Top2 proteins

For biochemical characterization of Top2-RG and Top2-FY,RG, the relevant mutations were introduced into plasmid pGAL1Top2 (37). The proteins were then overexpressed in and purified from a *top1*Δ background in order to eliminate the confounding effects of Top1 activity. The relaxation activities of the purified proteins were assessed using negatively supercoiled pUC18 DNA as substrate (38). Under our standard conditions, complete relaxation of pUC18 by the WT or Top2 mutant proteins required 25-50 ng of protein (Figure 2A). This result indicates that the overall catalytic activity of the mutant proteins was similar to that of WT yeast Top2.

**Figure 2.**
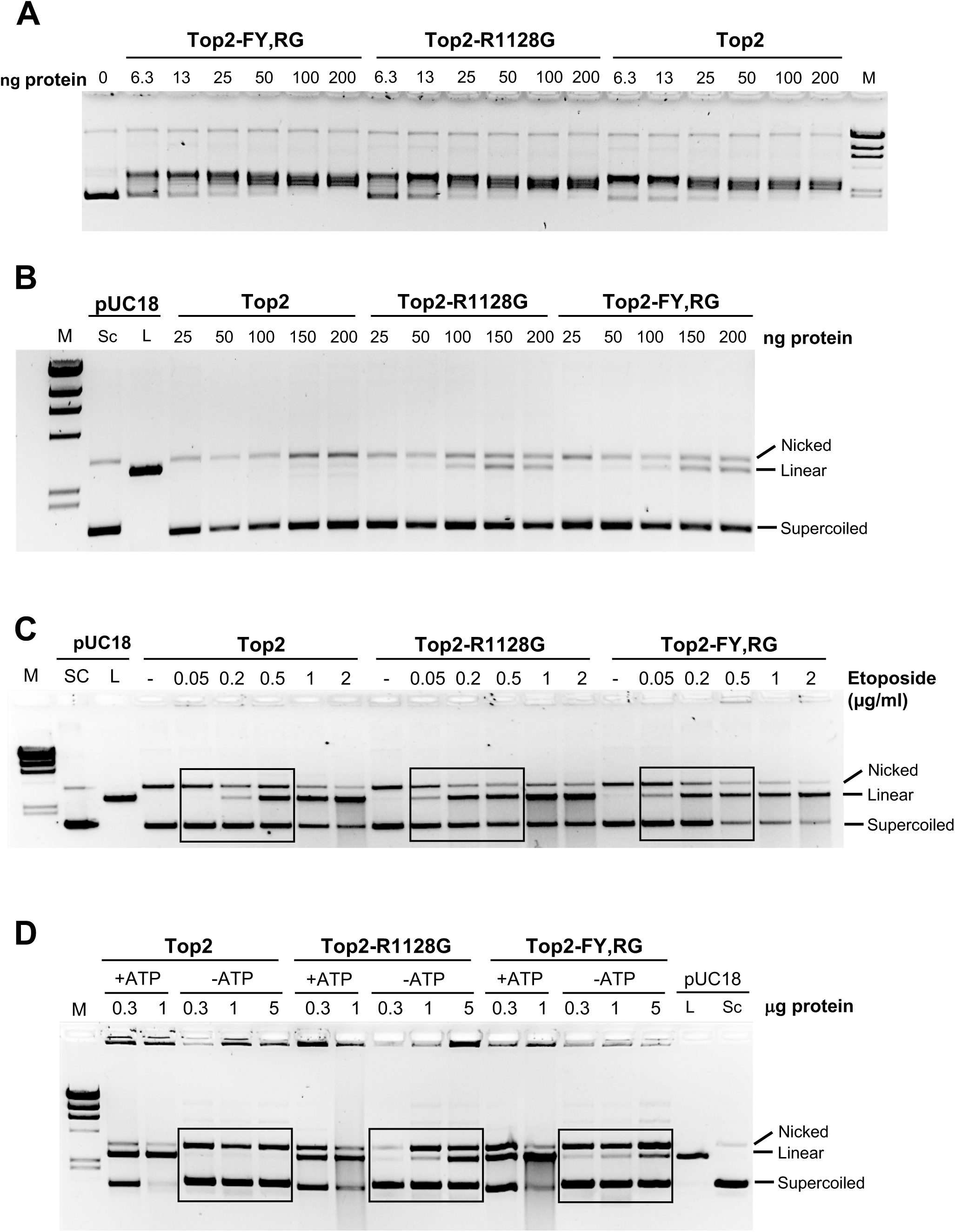
Biochemical activities of WT and mutant proteins. For purification, Top2 proteins were expressed from the *pGAL1* promoter in the *top1*Δ strain JELt1 (29). (A) DNA strand-passage activity of purified Top2-R1128G and Top2-FY,RG proteins was compared to the WT enzyme using 200 ng of negatively supercoiled pUC18 DNA; the amount of enzyme is indicated above each lane. (B) Top2 mutant protein mediated DNA cleavage of 200 ng negatively supercoiled plasmid DNA (pUC18) compared to WT enzyme. (C) Top2 mutant protein mediated DNA cleavage of plasmid DNA compared to WT enzyme in the presence of etoposide. (D) Top2 mutant protein mediated DNA cleavage of plasmid DNA compared to WT enzyme in the presence/absence ATP. Data in the presence of mAMSA are in Figure S1.

We next examined the ability of the mutant Top2 proteins to cleave pUC18 DNA in the absence of Top2 targeting agents. Double-strand DNA cleavage of plasmid DNA results in the formation of linear DNA molecules. For WT Top2 a very faint linear band was seen with 150 ng of purified protein (Figure 2B). By contrast, linear fragments were observed with Top2-RG and Top2-FY,RG at protein concentrations as low as 50 ng. We also examined the response of Top2-RG and Top2-FY,RG to etoposide (Figure 2C) and mAMSA (Supplemental Figure S1). In the presence of these small-molecule inhibitors, the mutant proteins were associated with a higher level of linear DNA as the drug concentration was increased. At higher etoposide concentrations (e.g., Top2-RG at etoposide concentrations greater than 1 μg/ml) smearing of DNA was observed, which presumably represents plasmids cleaved at two or more separate sites. These results demonstrate that the purified Top2-RG and Top2-FY,RG proteins have an intrinsic hypersensitivity to mAMSA and etoposide that is in agreement with the phenotypes of Top2-RG and Top2-FY,RG *in vivo*.

Although ATP is required for supercoil relaxation and for decatenation by Top2, eukaryotic Top2 cleaves DNA, albeit at a much lower level, even in the absence of ATP (3). To further examine potential mechanisms of Top2 trapping, we examined stable cleavage of pUC18 DNA by the WT and mutant proteins in the absence of ATP (Figure 2D). For WT Top2, linearization of pUC18 was barely detectable, although DNA nicking was seen. By contrast, linearized DNA was readily detected when pUC18 was incubated with 1 μg Top2-RG or 0.3 μg Top2-FY,RG. These results demonstrate that ATP is not required for elevated DNA cleavage by the mutant proteins, although it still strongly potentiates cleavage. Because ATP (or a non-hydrolyzable analog) is required for progression through the catalytic cycle and for strand passage, these results suggest that progression of the catalytic cycle is not required for elevated, drug-independent cleavage of DNA by these proteins.

### Top2-FY,RG is mutagenic and is associated with *de novo* duplications

Expression of the *top2-FY,RG* allele resulted in a hyper-recombination phenotype and was lethal in a *rad52*Δ background (Figure 1), consistent with formation of potentially toxic DSBs. *In vitro*, the mutant protein generated persistent nicks as well as DSBs (Figure 2), leading us to examine whether its expression might be mutagenic. For this analysis the EV, pDEDTop2 or pDED1Top2-FY,RG was introduced into a haploid *TOP2* background. The *CAN1* forward-mutation assay was used to measure mutation rates and analyze mutation types. In this assay any mutation that disables function of the encoded protein confers resistance to canavanine, a toxic arginine analog. The canavanine-resistance (Can-R) rate was indistinguishable in cells containing either the EV or pDED1ToP2. By contrast, expression of the *top2-FY,RG* allele elevated the Can-R rate 2.7-fold (Table S1).

Approximately 75% of *can1* mutations detected were base substitutions when either the EV or the pDED1Top2 plasmid was present (105/142 and 111/155, respectively; p=0.74 by contingency Chi-square; see Table S2 for complete spectra). By contrast, only 28% of mutations were base substitutions in the strain containing the pDED1Top2-FY,RG plasmid (50/176; p<0.0001). The proportional decrease in base substitutions was accompanied by a large increase in insertions of more than one base pair (from 2/142 with the EV to 75/176; p<0.0001). Most of these insertions (49/75) corresponded to *de novo* duplications, which are defined as the creation of a repeat where one did not previously exist. In the segment of the *CAN1* ORF shown in Figure 3A, for example, CTGT (nt 1239-1242) became CTGT<u>CTGT</u> in one mutant and CATT (nt 1272-1275) was duplicated to CATT<u>CATT</u> in another. The remainder of insertions >1-bp occurred within a pre-existing repeat; because of the genetic requirements for their formation (see below), we consider these jointly with the *de novo* duplications. The most frequent size of duplications was 4 bp (Table S2), which matches the distance between Top2- generated nicks and is relevant to the proposed mechanism of duplication formation (see Discussion).

**Figure 3.**
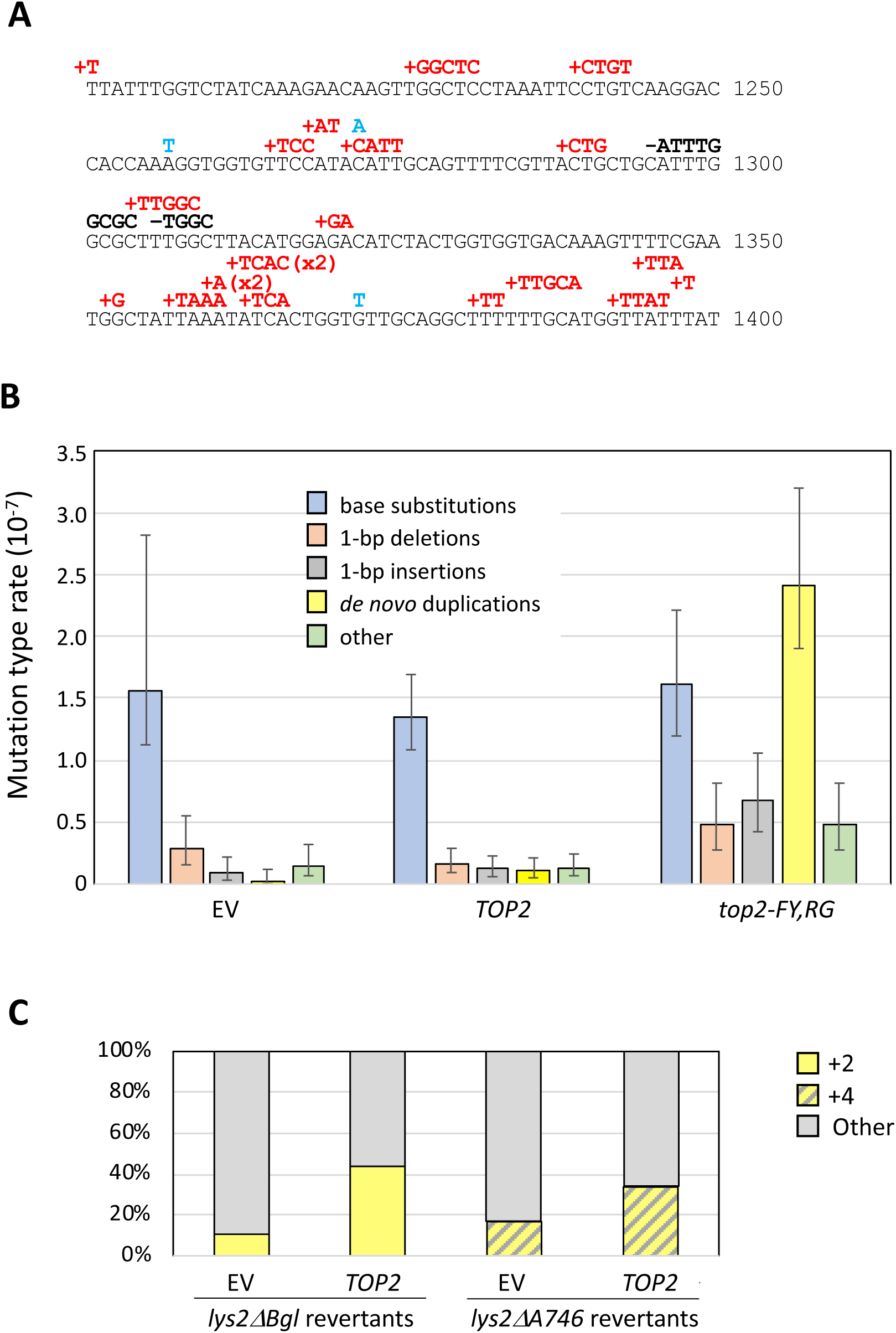
Top2-FY,RG expression is associated with *de novo* duplications. (A) Partial *CAN1* mutation spectrum. Mutations are above the sequence; insertions are in red, base substitutions in blue and deletions in black font. (B) Rates of specific mutation types in a *TOP2* strain containing the EV, pDED1Top2 or pDED1Top2-FY,RG. Error bars are 95% confidence intervals. (C) Proportions of 2-bp and 4-bp duplications among Lys^+^ revertants of the *lys2*Δ*Bgl,NR* (SJR1467) and *lys2*Δ*A746,NR* (SJR1468) alleles (28). Rates and associated spectra are in Tables S1-S2. Complete Lys^+^ spectra are in Figure S2.

The rates of specific mutation types (base substitutions, 1-bp deletions, 1-bp insertions, *de novo* duplications and others) in the presence of the EV, the pDED1Top2 plasmid or the pDED1Top2-FY,RG plasmid are presented in Figure 3B. We estimate an ∼80-fold increase in the rate of duplications, but no change in the base substitution rate, when the *top2-FY,RG* allele was overexpressed. It should be noted that there was also a 7.6-fold increase in the rate of +1 insertions associated with pDED1Top2-FY,RG. Although most of these 1-bp insertions were in short homopolymer runs and could reflect DNA polymerase slippage during replication, their increase suggests that many were likely generated by the same mechanism as the larger insertions.

### Top2 overexpression elevates duplications in frameshift reversion assays

Although there was no increase in the overall rate of Can-R in the presence of the pDED1Top2 plasmid, there was an ∼4-fold increase in duplications. Given their small number, however, neither the proportional increase in these events (2/142 for EV and 9/155 for pDED1Top2; p=0.09) nor their corresponding rate was significantly increased relative to cells that did not overexpress Top2. To focus specifically on 2- and 4-bp duplications, we used frameshift-reversion assays where similar events were previously observed (28). The *lys2*Δ*Bgl,NR* allele reverts by acquisition of a net -1 frameshift and so can detect 2-bp insertions; the *lys2*Δ*A746,NR* reverts by net +1 frameshifts, which includes 4-bp insertions. Top2 over-expression was accompanied by a proportional increase in 2-bp (from 3/28 to 22/50; p=0.0056) as well as 4-bp duplications (from 6/36 to 33/98, p=0.088; Figure 3C and Figure S2). When these events were jointly considered, their increase was highly significant (p=0.0014), demonstrating an association of a specific class of mutations with Top2 overexpression.

### Top2cc-dependent duplications require the NHEJ pathway

*In vitro*, the Top2-FY,RG protein generates nicks in addition to DSBs (Figure 2), either of which potentially could initiate the *de novo* sequence duplications observed *in vivo*. Because the NHEJ pathway can introduce sequence changes at the junction of joined ends and was previously implicated in generating the *de novo* duplications in the frameshift-reversion assays described above (28), we examined the relevance of this pathway to the mutagenesis associated with stabilization of the Top2 cleavage complex (Top2cc; Figure 4A and Tables S1-S2). We first deleted the *DNL4* gene, which encodes the DNA ligase required for NHEJ (39, 40), in cells containing the pDED1Top2-FY,RG plasmid. In the *dnl4*Δ background, there was a 40% reduction in the Can-R rate that was accompanied by a large proportional decrease in *de novo* duplications (from 75/176 to 7/194; p<0.0001) as well as 1-bp insertions (from 21/176 to 7/194; p=0.005). The Ku complex is also required for NHEJ in yeast, and results in *ku70*Δ background were similar to those obtained in the *dnl4*Δ background (Table S1). In addition to the requirement for Dnl4 and Ku, most end/gap-filling that occurs during NHEJ requires DNA polymerase 4 (Pol4). Deletion of *POL4* from the strain containing the pDED1Top2-FY,RG plasmid also significantly reduced the Can-R rate and the proportion of duplications (from 75/176 to 25/126; p<0.0001). The effect of *POL4* loss on duplications, however, was not as dramatic as that observed in the absence of *DNL4* (25/126 and 7/194, respectively; p<0.0001). Loss of *POL4* had no effect on 1-bp insertions (21/176 and 12/126 in WT and *pol4*Δ, respectively; p=0.63), which is consistent with its reduced requirement for duplications and the relatively high background of +1 events. Together, these data demonstrate that the majority of sequence duplications in this system are products of Top2-generated DSBs that are repaired by NHEJ.

**Figure 4.**
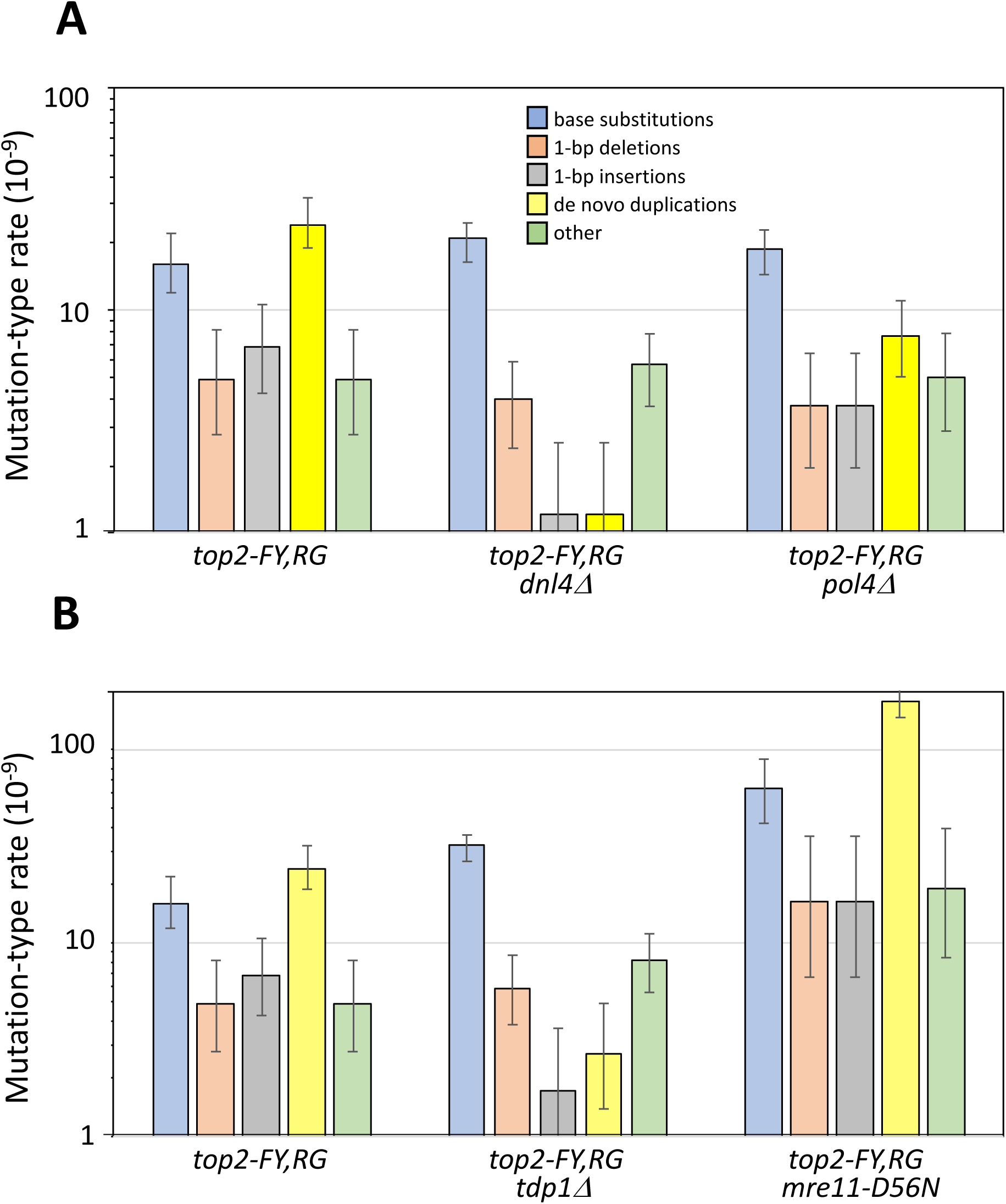
Genetic control of mutations associated with *top2-FY,RG* expression. Indicated candidate genes were deleted in the *TOP2* background and the resulting strains were transformed with EV, pDED1Top2 or pDED1Top2-FY,RG. (A) Duplications require Dnl4, and hence the NHEJ pathway, and partially require Pol4. (B) Duplications are promoted by Tdp1 and suppressed by the nuclease activity of Mre11 (*mre11-D56N*). Rates and associated spectra are in Tables S1-S2.

### Pathways for Top2cc removal affect duplication rates

A stabilized yeast Top2cc can potentially be removed by either (1) proteolytic degradation followed by peptide extraction through cleavage of the phosphotyrosyl bond or (2) nuclease-dependent removal of a Top2-linked oligonucleotide. In the first pathway, Top2 is presumably degraded by the proteasome and the remaining peptide removed by tyrosyl-DNA phosphodiesterase 1 (Tdp1; (41)). Although *TDP1* loss had no significant effect on the Can-R rate in the presence of the pDED1Top2-FY,RG plasmid, there was a large proportional reduction in duplications (75/176 to 11/205; p<0.0001) as well as 1-bp insertions (21/176 to 7/205; p=0.003) and rates of these events were significantly reduced (Figure 4B and Tables S1-S2).

In higher eukaryotes, a major pathway for Top2cc removal from a DNA end is by the endonuclease activity of the MRE11 component of the MRN (MRE11-RAD50-NBS1) complex, which releases a Top2-linked oligonucleotide (42). In *S. cerevisiae*, MRX (Mre11-Rad50-Xrs2) is the equivalent complex, and loss of any of the component proteins confers etoposide hypersensitivity (43). Although we were unable to obtain viable colonies following transformation of *mre11*Δ cells with the pDED1Top2-FY,RG plasmid, transformants were readily obtained in a background containing the nuclease-dead *mre11-D56N* allele (44). The essential role of yeast MRX in dealing with Top2-associated damage is thus distinct from its nuclease activity. Expression of the *top2-FY,RG* allele in an *mre11-D56N* allele background was associated with a 7.3-fold increase in the rate of *de novo* duplications (Figure 4B), indicating that Mre11 nuclease activity is primarily responsible for yeast Top1cc removal from DNA ends. In its absence, the phosphotyrosyl peptide that remains after Top2 proteolysis is cleanly removed by Tdp1 and the ends give rise to NHEJ-dependent duplications.

### Mutagenic effects of etoposide

To further examine the dependence of duplications on Top2cc stabilization, we isolated Can-R mutants in a drug-sensitized *TOP2* strain grown in the presence of either 200 μg/ml etoposide or DMSO, the vehicle for etoposide. Although etoposide did not alter the median Can- R frequency (Table S3), there were significant changes in the corresponding spectrum and in the frequencies of some mutation types (Figure 5). Relative to the DMSO-treated cultures, there was a proportional increase in duplications (from 1/162 to 36/222; p<0.0001) as well as 1-bp insertions (from 3/162 to 19/222; p<0.0001) in the presence of etoposide. Confirmation that the mutagenic effect of etoposide was mediated through Top2cc stabilization was obtained using the etoposide resistant *top2-5* allele (45). Because this allele confers temperature sensitivity, experiments were performed at room temperature (RT) rather than 30°. Interestingly, the Can-R frequency in the WT strain was elevated ∼2-fold when cells were grown at RT in the presence of DMSO or etoposide (Figure 5 and Table S3). As at 30°, however, etoposide treatment of the WT strain at RT stimulated duplications (from 4/288 to 18/305; p=0.007 by Chi-square) as well as 1- bp insertions (9/288 and 27/305; p=0.006), and their rates were not significantly different from those observed at 30°. By contrast, etoposide stimulated neither duplications (5/344 in DMSO and 4/297 in etoposide, respectively; p=1 by Fisher 2-tailed) nor 1-bp insertions (8/344 and 7/297; p=1) in the *top2-5* background.

**Figure 5.**
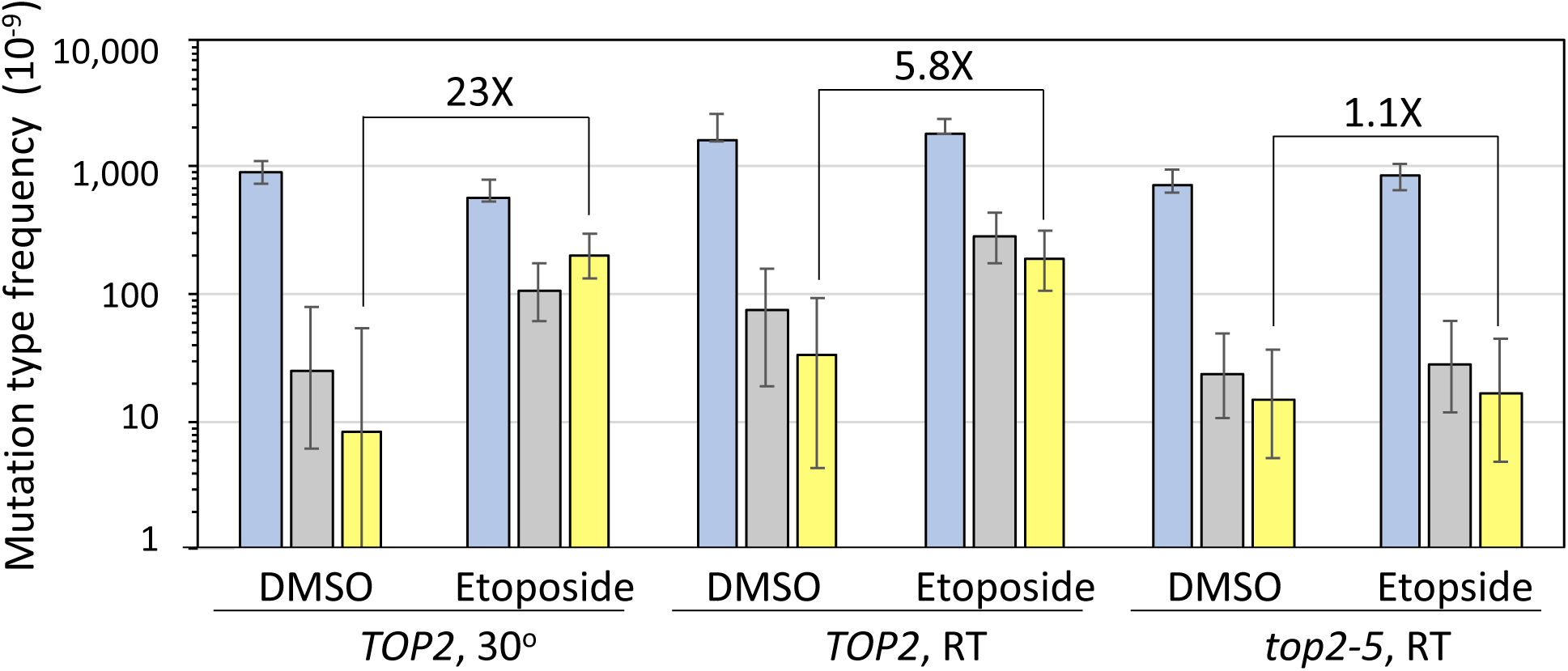
Etoposide stimulates duplications through its interaction with Top2. A *TOP2 pdr1DBD-CYC8::LEU2* (drug-sensitized) strain was grown in the presence of DMSO or DMSO+etoposide at RT or at 30°. The temperature-sensitive *top2-5 pdr1DBD-CYC8::LEU2* strain only grows at RT and is etoposide resistant. Median Can-R frequencies were determined rather than rates because of the variation in the number of viable cells in etoposide-treated cultures; error bars are 95% confidence intervals. Median frequencies and associated spectra are in Tables S2-S3.

## DISCUSSION

Topoisomerases carry out essential reactions that require DNA cleavage and a failure to quickly re-ligate DNA can lead to genome destabilization (21). In the current study, we identified and characterized yeast *top2* alleles (*top2-R1128G* and *top2-F1025Y,R1128G*) that produce proteins that are defective in quickly following up cleavage with re-ligation. Although the mutant proteins supported the essential function of Top2 *in vivo*, they conferred lethality in strains defective in the recombinational repair of DSBs or in the MRX complex. Consistent with elevated DSBs, overexpression of the Top2-FY,RG protein conferred hypersensitivity to Top2 poisons, was associated with a mitotic hyper-recombination phenotype and led to elevated levels of covalent Top2-DNA complexes. All of these phenotypes are consistent with the elevated DNA cleavage observed biochemically and are similar to the effects of etoposide in WT cells.

Although the Top2-FY,RG protein shows elevated cleavage, both amino acid substitutions map to the C-terminal dimer interface or ‘C-gate’ region (Figure 6A) rather than near the catalytic tyrosine. The C-gate constitutes the primary dimer interface that holds Top2 subunits together in the absence of DNA or ATP (46-48). During the Top2 catalytic cycle, this interface splits apart to allow a newly transported duplex that has crossed the cleaved-DNA segment to exit the enzyme (49). The stability of the C-gate interface, together with that of the nucleotide-dimerized ATPase domains, is thought to hold the two Top2 subunits together and guard against the accidental dissociation of the dimer during its duplex cleavage and passage reaction. Such dissociation would be expected to lead to persistent DSB formation.

**Figure 6.**
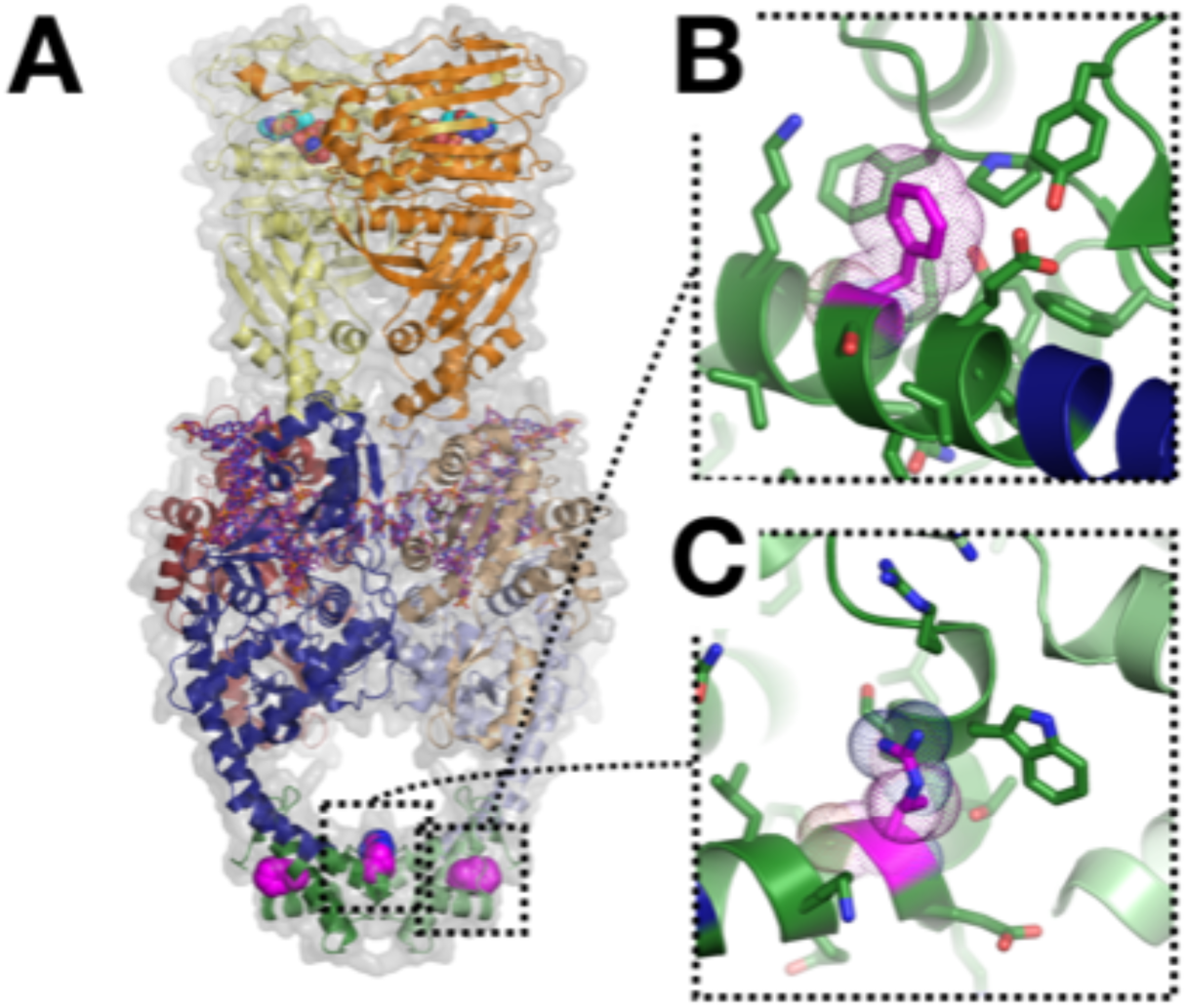
The F1025Y/R1128G mutations map to the Top2 C-gate. (A) Cartoon representation of a full-length, nucleotide-bound *S. cerevisiae* Top2 dimer bound to a cleaved DNA segment (PDB ID 4GFH; {Schmidt, 2012, Structure of a topoisomerase II-DNA-nucleotide complex reveals a new control mechanism for ATPase activity}). One subunit is colored in dark hues and the other in light. Dark/light-orange – ATPase domains; dark/light-red – TOPRIM Mg2+-binding domain; dark/light-blue – principal DNA binding region; dark/light-green – C-gate; dark-purple – cleaved DNA. AMPPNP is shown as cyan spheres; the locations of F1025 and R1128 are marked as magenta spheres. (B) Close-up of the region around F1025 is highlighted, with F1025 shown in magenta. (C) Close-up of the region around R1128, illustrating the packing of this residue adjacent to W1122 at the dimer interface.

A close up of the interface shows that F1025 protrudes from the coiled-coil arms of the principal DNA-binding region and nestles into a small hydrophobic pocket on one side of the globular region of the C-gate (Figure 6B). Given the location of F1025 and the protrusion of the tip of its benzyl ring so that it is solvent exposed, it seems unlikely that its replacement with tyrosine, which would project its phenolic oxygen into solution, would have an effect on C-gate stability. By comparison, R1128 packs against a tryptophan (W1122) that directly forms part of the dimer interface (Figure 6C). This interaction, together with the introduction of a glycine in the middle of a helical element, would be expected to locally destabilize the region. This could impact the integrity of the C-gate or the kinetics with which the dimer interface separates and re-associates. Either behavior could, in turn, detrimentally affect the cleavage propensity of Top2 and increase the life-time of the cleaved DNA state, the possibility of subunit dissociation, or both.

The hyper-recombination phenotype associated with Top2-FY,RG expression in yeast, as well as the synthetic lethality in the absence of recombination, are consistent with stabilization of the cleavage intermediate. A mutator phenotype was not expected, however, and could reflect either the persistence of nicked intermediates (see Figure 2) or errors associated with the repair of DSBs. The sequencing of Can-R mutants was particularly informative, revealing a shift from predominantly base substitutions to a rarely observed type of insertion: *de novo* duplication of 2- to 4 bp of sequence. The largest class of duplications was 4-bp in size, which matches the distance between the nicks Top2 makes on complementary DNA strands, and all duplications were NHEJ dependent. As illustrated on the left side of Figure 7, the complete filling in of the 5’ overhangs generated by Top2 cleavage, followed by NHEJ-mediated ligation, creates a 4-bp duplication.

**Figure 7.**
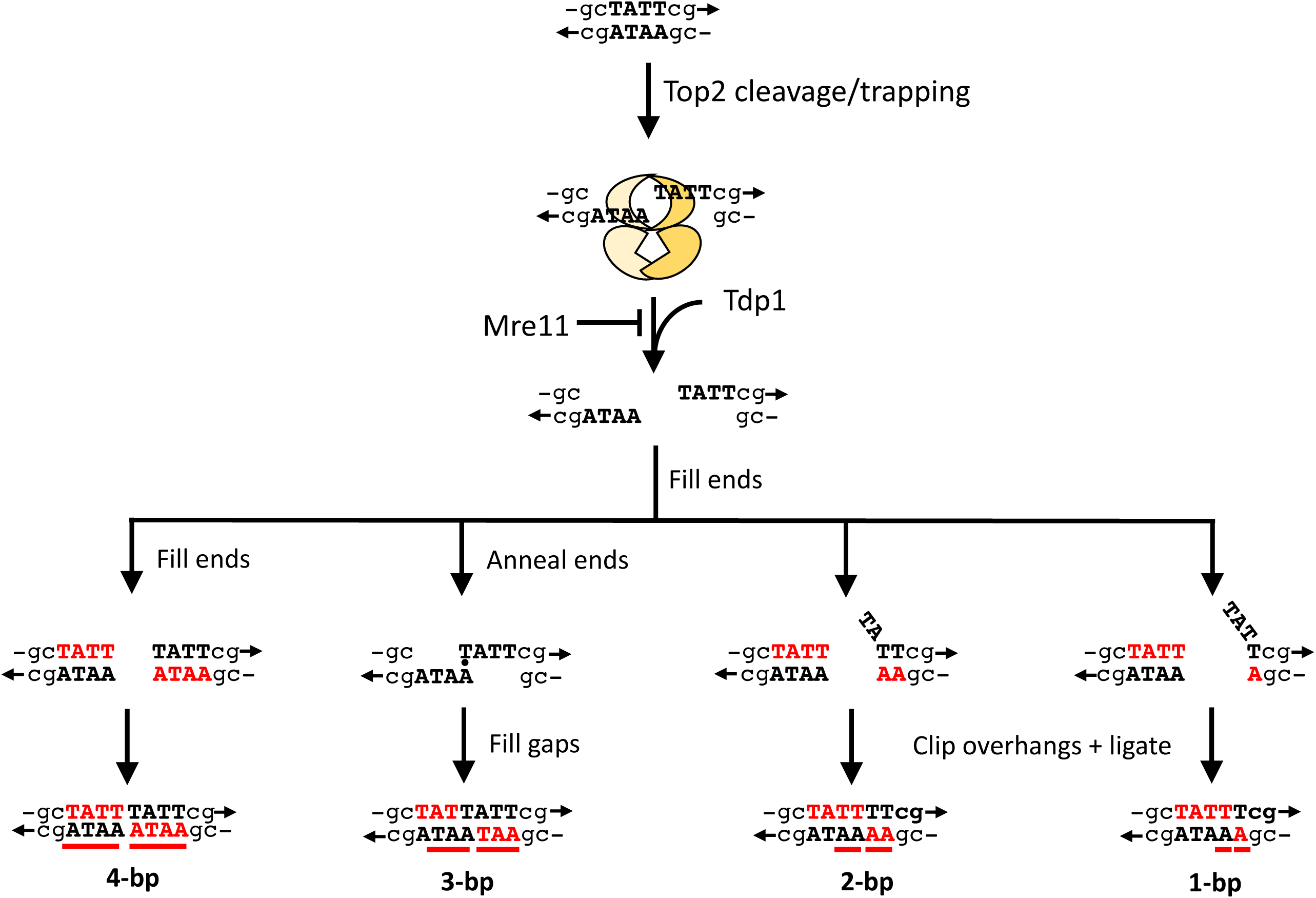
Models for Top2-associated deletions. Complementary strands are shown with arrows indicating 3’ ends; the 4 bp flanked by Top2 nicks is in bold capital letters. Each Top1 monomer (different shades of yellow) uses an active-site tyrosine to nick one DNA strand, forming a covalent phosphotyrosyl bond with the 5’ end and creating a recessed 3’-OH end. Mre11 suppresses duplications by nicking the 5’-terminal strand and thereby removing Top2 as part of a long oligonucleotide. Tdp1 cleaves the phosphotyrosyl bond to create a clean 5’ end. New bases added to recess 3’ ends after Top2 removal are in bold red font and duplicated segments are underlined in red. There are multiple ways to generate 1- to 3-bp duplications that depend on where Top2 cleaves the duplex and whether/how ends anneal before fill-in reactions.

A composite spectrum of insertions >1 bp that were identified in the WT and *mre11- D56N* backgrounds (Figure S3) suggests the occurrence duplication hotspots. These could reflect sites of preferred Top2-FY,RG cleavage/stabilization or sites where end filling is more frequent and NHEJ is relatively error prone. The positions of Top2 cleavage can be inferred from 4-bp duplications, but not for smaller duplications. These cleavage sites were aligned and analyzed for possible sequence conservation, such as the weak dyad symmetry reported for human TOP2 cleavage *in vitro* (50). Although none was observed in the nick-flanking region where the Top2 monomers are expected to contact DNA, there was a strong preference for AT base pairs in the region between the nicks (Figure S4). We suggest that the AT-richness may increase the stability of the cleavage intermediate or facilitate the strand separation that precedes the filling of 3’-recessed ends. Although stronger dyad symmetry has been associated with etoposide/VP16-stabilized human TOP2 and yeast Top2 cleavage intermediates (50, 51), there were not enough duplications identified following etoposide treatment to perform a similar analysis in the current study.

The Top2 cleavage site depicted in Figure 7 corresponds to position 1007, where ∼10% of the 4-bp duplications occurred (Figure S3). This position was also one of the few sites where 3-bp duplications were detected. The rarity of 3-bp duplications presumably reflects the fact that, unlike the 2- or 4-bp insertions, the addition of 3 bp maintains the correct reading frame of *CAN1* and rarely disables protein function. As illustrated in Figure 7, 3-bp duplications can be readily formed at position 1007 by annealing the terminal nucleotide at each end of the TATT cleavage site and then filling the adjacent gaps. Three-bp duplications could also arise by removing a single nt from one end of the DSB, followed by end filling and ligation. As illustrated in Figure 7, 2-bp *de novo* duplications require the addition of 4-nt to one end and 2-nt to the other; however, if a Top2 cleavage site encompasses a dinucleotide repeat, misaligned pairing between complementary strands would seem the more likely mechanism. Finally, 1-bp insertions can be generated by partial filling of one or both ends, or by misaligned pairing between the ends and subsequent gap filling. Based on the biochemistry of Top2, duplication sizes should not exceed the distance between the enzyme-generated nicks, and yet 21 of 163 (∼13%) of the duplications were 5 bp. Eight of these occurred at a single position that increased a 5-bp repeat (GGGCT) from two copies to three and most likely reflected DNA polymerase slippage. The remainder were not in repetitive sequence and raise the intriguing possibility that Top2 monomers might occasionally create nicks that are 5 bp instead of 4 bp apart. This may be a specific feature associated with the *top2-FY,RG* allele or other mutations that destabilize the dimer interface.

Before end-filling and ligation can occur, the trapped Top2 must be removed from DNA ends. The increase in *de novo* duplications observed in the absence of the Mre11 nuclease activity (*mre11-D56N* background) indicates that, as in higher eukaryotes (42), Top2 is removed by the MRX-Sae2 complex (MRN-CtIP in mammals). In contrast to mammalian cells, however, the nuclease-dead Mre11 protein was compatible with viability; this may simply reflect a much lower load of persistent Top2 damage in the much smaller yeast genome or a more robust role of yeast Tdp1 in Top2cc removal from DNA ends. Although Mre11 nuclease activity was not required for viability in the presence of overexpressed Top2-FY,RG, it was necessary for the MRX complex to be present. This requirement does not reflect its essential role in yeast NHEJ, as NHEJ was dispensable for survival. In this context, the MRX complex may be required for mediating an appropriate checkpoint response (52).

*In vitro*, Mre11 nicks 15-40 nt from a 5’-blocked end (53), which precludes the end filling required for *de novo* duplications. 5’-end cleavage also facilitates more extensive end resection and commits repair to HR (54), although microhomology-mediated end joining (MMEJ) is an alternative outcome if HR is not possible (55). It should be noted that the mutation assay done here was in haploid cells, where the only recombination option was the identical sister chromatid. The dramatic increase in *de novo* duplications in the absence of Mre11 nucleolytic activity indicates that MRX is the primary remover of trapped Top2, as it is in mammalian cells. Loss of Tdp1 had the reverse effect and duplications were virtually eliminated. Although Tdp1 was originally identified as a phosphodiesterase that removes 3’-linked peptides from DNA (56), it also has been implicated in removal of 5’-linked peptides and its loss confers etoposide sensitivity (41). It should be noted that yeast does not have a protein analogous to the TDP2 protein of mammals, which also is important for TOP2 removal (57) and suppresses chromosome rearrangements by creating DSB termini that are substrates for NHEJ (58).

The broader implications of the Top2-dependent mutation signature described here are two-fold and derive from the highly conserved biochemistry of Top2 and subsequent repair mechanisms. First, the stabilization of the covalent-cleavage intermediate by chemotherapeutic drugs, or the presence of an appropriate mutant TOP2 protein, is expected to have a similar consequence in mammalian cells. Insertions are much more likely to disrupt gene function than are base substitutions, with even a very low level having detrimental consequences. This may contribute to secondary malignancies that arise following the clinical use of TOP2 inhibitors and the corresponding genomes would be expected to have a distinctive mutation signature. Mutations in TOP2 or protein overproduction could also be potential drivers of tumorigenesis. In this regard, we note that the yeast Top2 signature identified here matches insertion-deletion signature 17 (ID17) in human cancers (59). A second implication of Top2-associated mutagenesis is that it provides a mechanism for the birth of repetitive sequences, as demonstrated here in yeast strains that overproduce the WT protein. In addition to its essential role during genome duplication, Top2 activity may thus be an important contributor to genome evolution.

## ACKNOWLEDGEMENTS

We thank Steven Rozen (Duke-NUS) for pointing out that the yeast Top2 signature matches ID17 in COSMIC (Catalog of Somatic Mutations in Cancer). Research was supported by grants from the National Institutes of Health: R35 GM203587 to S.J-R, R03 CA216010 to J.L.N., and R01 CA077373 to J.B.

## MATERIALS AND METHODS

### Media and growth conditions

YPD (1% yeast extract, 2% Bacto-peptone, 2% dextrose, 250 mg/liter adenine; 2% agar for plates) was used for non-selective growth. Synthetic complete (SC) medium contained 0.15% yeast nitrogen base, 0.5% ammonium sulfate and 2% dextrose (2% agar added for plates) and was supplemented with all amino acids plus adenine and uracil. Drop-out plates missing one amino acid or base (e.g., SC-Ura medium contained no uracil) were used for selective growth. For the mitotic recombination experiments, diploid strains were grown in SC/peptone-Ura (SCP-Ura; 2% tryptone, 0.17% yeast nitrogen base lacking NH_4_SO_4_, 250 μg/ml adenine; 2% agar for plates). Canavanine-resistant (Can-R) mutants were selected on SC-Arg plates containing 60μg/ml L-canavanine sulfate. All growth at 30° unless otherwise indicated.

### Strain constructions

A list of all yeast strains is provided in Table S4. Haploid strains used for mutation analyses were *RAD5* derivatives of *W303* [*ade2-1 his3-11,15 ura3-1 leu2-3,112 trp1-1 can1-100 rad5-G535R*; (60)]. *DNL4, YKU70, POL4* or *TDP1* was deleted by one-step allele replacement using PCR fragments amplified from a plasmid containing a selectable drug resistance marker. The *natMX4* cassette was amplified from pAG25 (61) and *loxP-hph-loxP* from pSR955 (62). The nuclease-dead *mre11-D56N* allele was introduced using two-step allele replacement following transformation with the *Sph*I-digested pSM444 (63).

To increase intracellular accumulation of etoposide the *PDR1* gene, which regulates the expression of drug efflux proteins, was replaced with a *LEU2*-marked construct containing the DNA binding domain of Pdr1 fused to the repressor domain of Cyc8 (64). Prior to transformation with an appropriate *pdr1DBD-CYC8::LEU2* fragment, the *LEU2* locus was replaced with a *loxP- TRP1-loxP* cassette amplified from pSR954 (65) to prevent recombination with the introduced fragment. The *TRP1* marker was subsequently removed by expression of the Cre recombinase from plasmid pSH63 (66). The *top2-5* temperature-sensitive allele was introduced by two-step allele replacement following transformation with *Kpn*I-digested YIp5top2-5 (45) and selecting Ura^+^ colonies. Integration of this plasmid truncates the endogenous *TOP2* allele so that only *top2-5* is expressed.

### *In vivo* complex of enzyme (ICE) assay in yeast

A C-terminally, HA-tagged mutant Top2 was generated by amplifying a fragment of pDED1Top2*-*FY,RG containing the *top2-FY,RG* mutations (forward and reverse primers 5’- TGTCAACTGAACCGGTAAGCGCCTCTG and 5’-CTTTGTCTCCTTGATCGTTGTGGT, respectively) and the backbone of pDED1Top2WT-3HA (ref) containing the HA-tag (forward and reverse primers 5’-CCTAAATTGGCCAAGAAGCCAGTCAGGAA and 5’- AGCACCATAACCGTTTCTACCACCAGT). The two fragments were co-transformed into yeast and assembled by homologous recombination. pDED1Top2WT-3HA was constructed by replacing the 3’ end of the *TOP2* gene of pDED1Top2 with a *Kpn*I/*Sac*I fragment from pJBN214 (67) that contained Top2 tagged with 3xHA.

YMM10t2-4 containing the HA-tagged WT or mutant plasmid was inoculated into SC-Ura and incubated overnight at 34°. Cells were diluted to OD_600_=0.8 in 20 ml SC-Ura and after re-entering log phase (after approximately 2 h), DMSO or 20 μg/ml etoposide was added. After incubation for an additional 1 h at 34°, cells were harvested by centrifugation and washed in lysis buffer (6 M guanidinium thiocyanate, 1% sarkosyl, 4% Triton X-100, 1X TE, pH 7.5). A yeast protease inhibitor cocktail (Sigma-Aldrich; 50 μl per g of yeast cells) and dithiothreitol (DTT; final concentration 1%) were added prior to lysis. Cells were lysed at 4° using a Bead Beater (Biospec) for two 50-sec rounds. Lysates were incubated at 65° for 15 min, followed by a 15 min, 14,000 rpm centrifugation to remove cell debris. 400 μl of the lysates were mixed with 700 μl of 1% sarkosyl and the centrifugation was repeated. 1 ml of each lysate was mixed with 2 ml of 1% sarkosyl and then layered on 2 ml of 150% (w/v) cesium chloride (in 1X TE, pH 7.5) in 4.9 ml OptiSeal tubes (Beckman Coulter, cat #362185). Samples were centrifuged for 18 h at 42,000 rpm in an NVT 65.2 rotor (Beckman Coulter) at 25°. After removing the supernatant, pellets were washed with 500 μl of 95% ethanol and air-dried for 5 min. Pellets were resuspended in 200 μl of 1X TE and incubated at 65° for 5 min followed by a 1-h incubation at room temperature while shaking. RNase A was added to a final concentration of 100 μg/ml and samples were incubated for 1 h at 37°. Samples were ethanol precipitated, washed with 70% ethanol and air dried for 5 min. Pellets were dissolved in 50-100 μl of H_2_O, incubated at 65° for 5 min and then incubated for 1 h at room temperature incubation while shaking. DNA was quantitated using a UV spectrophotometer. 10 μg aliquots were digested with 0.5 μl micrococcal nuclease for 30 min at 37° micrococcal nuclease in 500 mM Tris-Cl pH 7.9, 50 mM CaCl_2_. The reaction was stopped by adding 4X Laemmli buffer, samples were boiled for 5 min and equal volumes were loaded onto a 4-15% gradient gel (Bio-Rad) for SDS-PAGE. After transfer to a PVDF membrane, Top2 covalent complexes were detected using an anti-HA (Santa Cruz, sc- 805) or anti-Top2 (TopoGen, #TG2014) antibody. In addition, 2 μg were transferred to nitrocellulose using a slot-blot apparatus (Bio-Rad) and an anti-dsDNA antibody (Abcam, #27156) was used to confirm equal amounts of DNA in each sample. After incubation with the corresponding secondary anti-rabbit antibody (GE Healthcare, NA934) or anti-mouse antibody (GE Healthcare, NA931), blots were visualized using SuperSignal™ West Femto ECL substrate (ThermoFisher Scientific, #34095).

### Western blotting

Yeast carrying either pDED1Top2-3HA or pDED1Top2*-*FY,RG-3HA were grown and diluted as described previously. Log-phase cells were pelleted and lysed using an alkaline lysis/glass bead procedure (68). Following neutralization, samples were digested with for 1 h on ice with micrococcal nuclease to release Top2 from DNA. Protein concentrations were determined using the Bradford assay (Bio-Rad). Lysates were mixed with 4X Laemmli buffer, boiled for 5 min, and 30 μg were loaded onto a 4-15% gradient gel (Bio-Rad) for SDS-PAGE. Separated proteins were transferred to a PVDF membrane and the membrane was incubated with antibodies to the HA epitope tag (Santa Cruz Biotechnology, sc-805) and α tubulin (Santa Cruz Biotechnology, sc-53030), followed by incubation with the corresponding secondary anti-rabbit antibody (GE Healthcare, NA934) or anti-rat antibody (Santa Cruz Biotechnology, sc- 2006). Blots were visualized as described above.

### Topoisomerase biochemical assays

Wild-type and mutant Top2 proteins were overexpressed from YEpGALyTop2 (69) in yeast strain JEL1t1 (29) and were purified as previously described (38). Top2 relaxation assays were carried out as previously described (38). Briefly, reactions were carried out in a final volume of 20 μl containing 50 mM Tris-HCl, pH-8.0, 100 mM KCl, 1 mM EDTA, 8 mM MgCl_2_, 2% glycerol, and 1 mM ATP; purified Top2 as indicated; and 200 ng of pUC18 DNA. pUC18 was purified from *E. coli* using a Qiagen plasmid Midi kit. After incubation at 30o for 15 min, reactions were quenched with 10mM EDTA and analyzed on 1% agarose gels in 1X TAE. Conditions for Top2 DNA cleavage reactions were the same as for Top2 relaxation assays, but reactions were quenched with SDS. Following treatment with proteinase K for 2 h at 50o, samples were extracted sequentially with phenol, phenol-chloroform (50:1) and chloroform. DNA was collected by ethanol precipitation and analyzed by agarose gel electrophoresis as above. pUC18 DNA linearized with *Eco*RI was used as a standard in cleavage assays.

### Mutation and recombination rate measurements

The *TOP2* or *top2-FY,RG* allele was constitutively expressed from the *DED1* promoter (*pDED1*) in a YCp50-derived (34) centromeric plasmid with *URA3* selectable marker (30). Following transformation, plasmid presence was selected and cells subsequently maintained on SC-Ura plates. For mutation rate measurements, individual colonies containing the EV, *TOP2*, or *top2-FY,RG* plasmid were inoculated into 1 ml of SC-Ura medium and grown to saturation (3 days) on a roller drum. Appropriate dilutions were plated on SC-Ura and SC-Arg+Can plates to determine the number of viable cells and Can-R mutants, respectively, in each culture. Colonies were counted after 3 days and mutation rates were calculated using the method of the median (70); 95% confidence intervals (CIs) were determined as previously described (71). Recombination rates were similarly measured by plating saturated cultures on the appropriate selective medium.

For the etoposide experiments, the *top2-5* mutant was grown at room temperature; the WT strain was grown at either 30° or at room temperature. A given experiment was begun by inoculating a single colony into 60 ml SC-complete liquid medium. The resulting culture was then divided and either etoposide (200 μg/ml dissolved in DMSO) or an equivalent volume of DMSO was added to each half. Each of these cultures was then further separated into individual 5-ml cultures, and these were incubated for 3 days on a roller drum. Cultures were washed two times re-suspended in 1 ml of H_2_O and each was sonicated prior to diluting and plating. Appropriate dilutions were plated on YPD and SC-Arg+Can as described above. Because of variation in the total number of cells in different experiments, median Can-R frequencies are reported; 95% CI for the median was determined as described above. Experiments for the *TOP2* strain were done at both room temperature and 30°, while the *top2-5* strain was grown only at room temperature.

### *CAN1* mutation spectra

Independent Can-R colonies were obtained using a pin-plating technique that generates ∼100 mini-cultures/plate. Cells were diluted in H_2_O and a 100-count, flat-tipped custom pinning device was used to transfer ∼10^3^ cells/pin onto SC-Ura plates. Cells were additionally spotted onto SC-Arg+Can plates to ensure that there were no pre-existing Can-R mutants. After 3 days of growth, cells were replica-plated onto SC-Arg+Can plates and were incubated for an additional 3 days. A similar technique was used to isolate independent Can-R mutants arising in the presence of 200 μg/ml etopside (control plates contained DMSO), with growth at 30° or room temperature, as appropriate. A single Can-R colony from each spot was used for genomic DNA extraction and subsequent sequencing.

The *CAN1* locus from genomic DNAs of mutants was amplified in a 96-well format using MyTaq DNA polymerase (Bioline). Each PCR product was uniquely barcoded using primers that contained 20 nt of *CAN1*-specific sequence conjugated to 16 nt of PacBio forward or reverse barcodes (https://github.com/PacificBiosciences/Bioinformatics-Training/blob/master/barcoding/pacbio_384_barcodes.fasta). The amount of each PCR product was estimated by running on an agarose gels and a similar concentration of each was used to construct the pool for subsequent SMRT sequencing. Following purification of the pooled DNA (GeneJet PCR Purification Kit, ThermoFisher Scientific), SMRT libraries were constructed and sequenced by the Duke Center for Genomic and Computational Biology or LabCorp (Burlington, NC) using the PacBio RSII and Sequel systems. Circular consensus sequence (CCS) reads were sorted by barcodes and analyzed using the in-house pipeline (SmrtSeqTool) previously described (72).

### *LYS2* Mutation Spectra

pDEDTop2 or EV was introduced into the *lys2ΔA746,NR* and *lys2ΔBgl,NR* strains SJR1467 and SJR1468, respectively (28). Transformants were selected and maintained SC-Ura plates. Independent Lys^+^ colonies were obtained using the pin-plating technique described above. After 3 days of growth, cells were replica-plated onto SC-Lys plates and incubated for an additional 5 days. A single Lys^+^ colony from each spot was used for genomic DNA extraction. The relevant portion of the *LYS2* gene was amplified by PCR and sequenced by GeneWiz (Durham, NC) using the primer 5’-GTAACCGGTGACGATGAT.

### Recombination experiments

The EV and pDEDtop2-FY,RG plasmids were transformed into a diploid yeast strain carrying multiple heteroallelic markers (*tyr1-1/tyr1-2, his7-2/his7-1, trp5-d/trp5-c* and *lys2-1/lys2- 2*). Cells containing the plasmids were selected on SC-Ura plates. Approximately 10^4^ cells were inoculated into 1 ml of SCP-Ura liquid medium and cultures were incubated for 4 days on a roller drum. Appropriate dilutions were plated on SCP-Ura plates to determine the number of viable cells in each culture and onto SC-Tyr, SC-His, SC-Trp, and SC-Lys plates to determine the numbers of recombinants. The median recombination frequency was reported for each heteroallelic marker because of the variation in total cell numbers in different experiments.

### Statistical methods

The proportions of mutation types in different strains were compared using a contingency Chi-Square or Fisher Exact test as appropriate (vassarstats.net); p<0.05 was considered significant. Rates were calculated using method of the median (70) and 95% confidence intervals (CIs) for rates and median frequencies were determined (73). Mutation-type rates or frequencies were calculated by multiplying the total Can-R rate or frequency by the proportion of the mutation type in the corresponding specturm. The 95% CI for each mutation type was determined by jointly considering the CI for the Can-R rate or frequency and the CI for the proportion (Vassarstats.net). This was done using the “root of the square of the sums” (RSS) or right triangle method (74). Rates/frequencies obtained in different strain backgrounds were considered significantly different if the respective 95% CIs did not overlap.

## SUPPLEMENTAL FIGURE LEGENDS

Figure S1. Effects of mAMSA on viability and Top2 cleavage activity. (A) Cell viability was determined by a yeast colony formation assay in JN362 *top2-4* cells carrying either pDED1Top2-F1025Y, pDED1Top2-R1128G, pDED1Top2-F1025Y,R1128G or pDED1Top2 (WT) plasmid after 24-hour exposure to mAMSA at 34°. Survival is relative to that at the time of mAMSA addition (100%). Yeast cells carrying the *top2*-*R1128G* allele were sensitive to mAMSA at concentrations as low as 5 µg/ml. (B) Plasmid cleavage assay with purified Top2 proteins. DNA cleavage of 200 ng of negatively supercoiled plasmid DNA (pUC18) was in the presence of mAMSA as indicated above each lane. Each enzyme produced DNA cleavage (linear and nicked products) in an mAMSA dose-response. At equal ng of purified protein, the mutant proteins demonstrated elevated levels of DNA cleavage in the absence of mAMSA as well as a greater response to mAMSA.

**Figure S2. Reversion spectra for the *lys2*Δ*A746,NR* -1 and *lys2*Δ*Bgl,NR* +1 frameshift alleles**. Strains contained either the EV or pDED1Top2 allele and the number (n) of Lys^+^ revertants sequenced from each is in parentheses. The positions of mutated homopolymer runs >3N are highlighted in yellow; the dotted vertical lines indicate the common sequence present in the reversion windows. +2 and +4 mutations are highlighted in gray in each spectrum.

**Figure 3. Composite *can1* spectrum of duplications >1 bp in WT and *mre11-D56N* strains containing pDED1-Top2-FY,RG**. Duplications >1 bp are positioned above the *CAN1* sequence (black). Those identified in the WT background are in bold and those from the *mre11-D56N* strain are in normal font; red corresponds to 2-4 bp duplications while blue indicates duplications >4 bp. The 5-bp insertions highlighted in gray are those occurring a pre-existing repeat (underlined).

**Figure S4. Sequence logo of 4-bp duplications in WT and *mre11-D56N* strains containing pDED1-Top2-FY,RG**. The logo was obtained by aligning the duplicated sequences (boxed) in 67 4-bp duplications (see Table S2). The lollipops indicate Top2 subunits, which are above/below the nucleotide to which each forms a covalent phosphotyrosyl link. By convention, the site of linkage on the top strand is labeled +1. The logo was constructed as described (76, 77) using website https://weblogo.berkeley.edu/.

